# Sparse-spectral microendoscopy for real-time visualization of tumor cell phenotype and microenvironment spatial heterogeneity *in vivo*

**DOI:** 10.1101/2022.06.17.496624

**Authors:** Bryan Q. Spring, Akilan Palanisami, Mohammad Ahsan Saad, Eric M. Kercher, Ryan T. Lang, Rebecca C. Harman, Jason Sutin, Zhiming Mai, Tayyaba Hasan

## Abstract

Cancer heterogeneity and its transformation with time propels treatment resistance and confounds patient outcomes. The inability to monitor *in vivo* the low abundance, heterocellular phenotypes that resist treatment and ultimately lead to patient death limits the ability to design precision therapies. Here we overcome limitations in multiplexed fluorescence phenotyping to introduce real-time, cellular resolution visualization of tumor heterogeneity *in vivo*. This method was performed to simultaneously map for the first time 5 individual biomarkers of stemness, proliferation, metabolism, leukocytes and angiogenesis deep within the peritoneal cavities of micrometastatic cancer mouse models at 17 frames per second (fps). The newly developed imaging system revealed distinct cancer cell phenotype–immune cell spatial correlations and clearly visualized the dynamic spatial response of resistant cancer cell niches following treatment. Furthermore, wide-field datasets were generated to facilitate derivation of a mathematical framework for quantifying biomarker spatial variation and thereby overcoming the area restrictions of conventional tumor biopsy. These results pave the way for real-time identification of cancer cell phenotypes in a clinical setting, on which optimized treatment regimens can be based for personalized treatment and precision therapy e.g., tumor margin determination during surgical resection. Additionally, this modality can be used to obtain more fundamental insights into tumor heterogeneity and how treatments affect the molecular and cellular responses of patient-specific disease.

## Introduction

Heterogeneity in malignant lesions, both in the cancer cell phenotypic expression and the tumor microenvironment, is a pivotal driver of treatment resistance and patient outcome variability^1^. Stem-like cell behavior^2^, crosstalk with immune cells^2^, proliferation status^1^ and metabolic activity^1^ have been implicated in treatment failure, and gross spatial variability has been observed for all of these factors. Sophisticated techniques for studying cancer cell heterogeneity, including *ex vivo* immunofluorescence^3,4^, imaging mass cytometry^5,6^ and barcoding^7^, have made tremendous strides in understanding the relationships between microscale tumor architecture and patient outcomes. Despite these advances, identifying the rare microenvironmental niches characterized by combinations of specific biomarkers that ultimately lead to treatment failure has proven challenging^8^. Fundamentally, the invasive nature of biopsy tissue sampling and its limited volume severely restricts the study of how different microenvironments respond to treatment *in vivo*– information necessary for developing therapies targeting resistant niches before they dominate the tumor biology. A single, elegant attempt at moving real-time phenotyping *in vivo* demonstrated the speeds necessary for intraoperative compatibility but has been unable to image more than 2 molecular markers^9^, which is insufficient for understanding the complex interplay between the immune system, stem-like cells, metabolism and tumor growth needed to rationally design combination therapy. A key limitation of these techniques arises from the quantum fluctuation (shot noise)–limited fluorescence signal that occurs at microscale resolution and real-time speed *in vivo*, leading to poor emission spectrum estimation.

## Results

To enable phenotypic mapping of the tumor microenvironment, we have developed “Sparse sensing of Multiplexed *In vivo* imaging with Real time Cellular resolution (SMIRC)”. SMIRC leverages properties of spectral sparseness (i.e., compressibility)^10^ to efficiently estimate pixel spectra from low abundance targets, enabling several molecular species to be simultaneously acquired and spectrally unmixed at real-time speeds using graphics processing unit (GPU)–accelerated algorithms (see Methods, Figures 1a–b and Supplementary Data Figs. 1–2). *In vitro* validation of fixed cancer cells clearly demonstrates cellular resolution imaging of five different markers, with the potential for increased marker detection by including additional near-infrared fluorophores (Supplementary Data Figs. 3–4).

**Figure 1.**
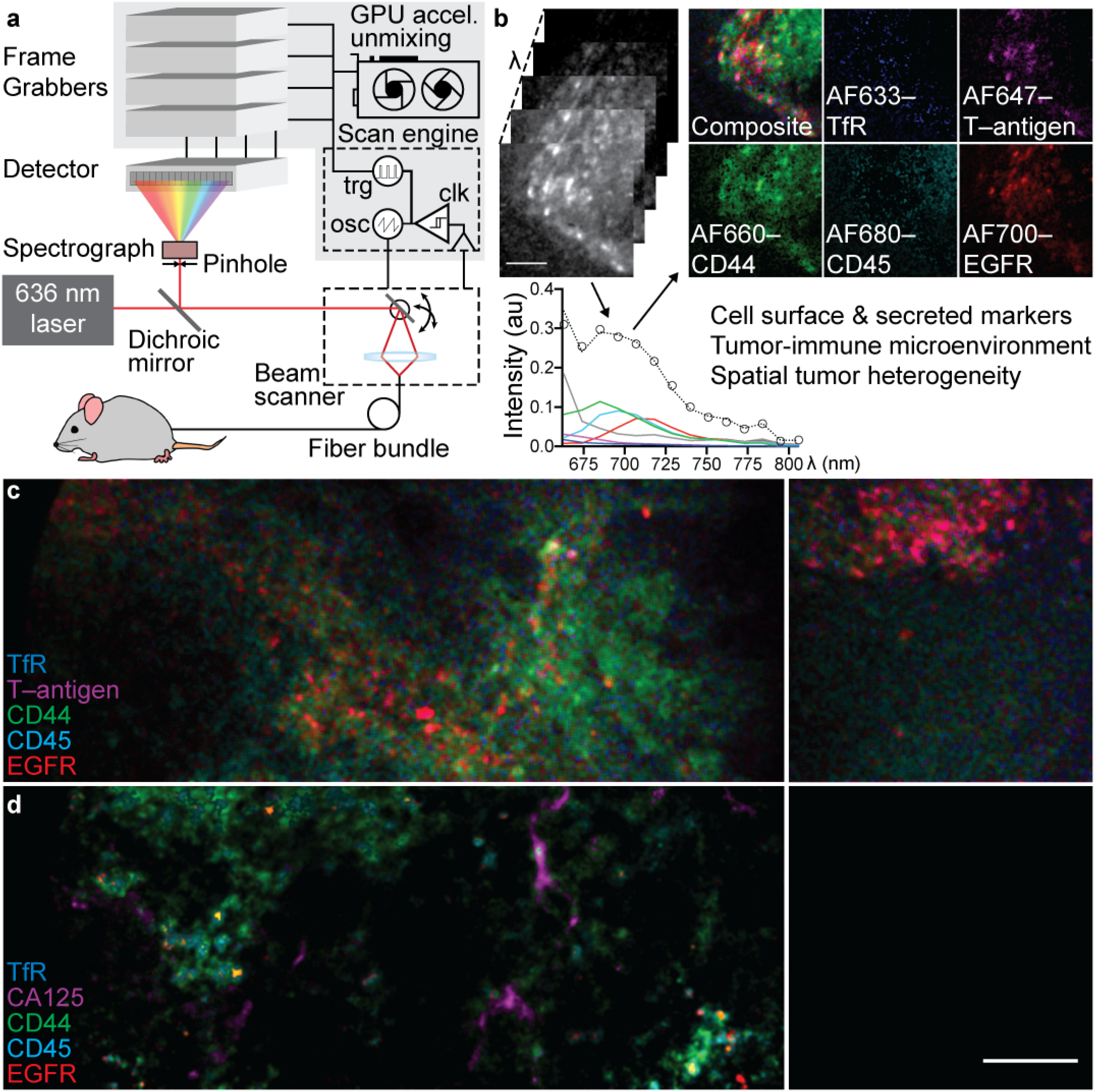
SMIRC overview. **a**, A functional schematic of the SMIRC instrumentation which outputs a stack of images spanning a range of emission wavelengths. **b**, The image stack is spectrally unmixed to recover abundance maps of fluorescent targeting conjugates, enabling unique spatiotemporal measurements of biomarker colocalization. Images taken from live mouse orthotopic pancreatic tumor. **c**, 5 target SMIRC of orthotopic pancreatic cancer (left panel) and a nearby metastasis on the peritoneal wall (right panel). **d**, 5 target SMIRC of peritoneally disseminated ovarian cancer (left panel) and tumor free control (right panel). All scale bars 100 µm. AF-Alexa Fluor; TfR-Transferrin

To demonstrate the ability of SMIRC to spatially characterize cancer-associated phenotypes *in vivo*, we imaged peritoneal disease in orthotopic mouse models of ovarian and pancreatic cancer (Figure 1c–d). Contrast agents were injected intraperitoneally 1 hour prior to imaging, and minimally invasive imaging was performed through a 1 mm incision via a catheter. Several well characterized molecular targets spanning a range of cancer hallmarks were imaged to illustrate the flexibility of multiplexed *in vivo* labeling including epidermal growth factor receptor (EGFR), CD44, CA125, transferrin uptake, Thomsen-Friedenreich carbohydrate antigen (T antigen) and CD45. EGFR has been validated in prior work as a sensitive *in vivo* cancer biomarker^11^ and is correlated with cell proliferation and survival. The cell surface glycoprotein CD44^12^ has been implicated in several reports as a biomarker for cancer stem-like behavior, mesenchymal phenotypes and chemoresistance. CA125 (MUC16)^13^ is currently the most reliable diagnostic marker for ovarian cancer and has been linked to immune modulation and metastasis. Transferrin uptake^14^ is an essential component of iron metabolism and is often elevated in cancer cells. The T antigen^15^ is a pan-carcinoma biomarker with suspected roles in proliferation and metastasis. Immune cell infiltration is a critical component of the tumor microenvironment and has significant prognostic value^16^. These effects are monitored here using CD45, a pan-leukocyte marker commonly used in flow cytometry. We routinely observed these and other biomarkers at the single cell level.

Spatial phenotypic architecture provides insight into tumor cell communication, microenvironmental niches and, crucially, the treatment response^17^, which provides guidance on the optimal timing of combination therapies. Existing real-time *in vivo* methods, based on label-free detection or conventional fluorescence methods, have been unable to resolve the phenotypic relationships that play a crucial role in cancer development. To extract quantitative phenotypic spatial correlations from SMIRC, we developed an automated phenotype identification and Euclidian distance mapping algorithm tailored to operate on the SMIRC datasets. The algorithm reduced the datasets into spatial maps of cancer cell subsets and immune cells (Figs. 2a–f) from which population and distance correlation metrics were evaluated (Figs. 2g–h). Chemoresistant, stem-like cancer cells^18^ were found to preferentially populate regions near immune cells^12,19^, in contrast to cancer cells with upregulated iron metabolism. These preferences were further exacerbated with sub-therapeutic photodynamic therapy (a cytotoxic light-based therapy)^20,21^. Post-treatment (30 hours), stem-like cancer cells came into intimate contact with a partner immune cell, demonstrating how dynamic changes in niche structure can be monitored and thereby used to inform the timing of combination therapies targeting changes in cellular communication.

**Figure 2.**
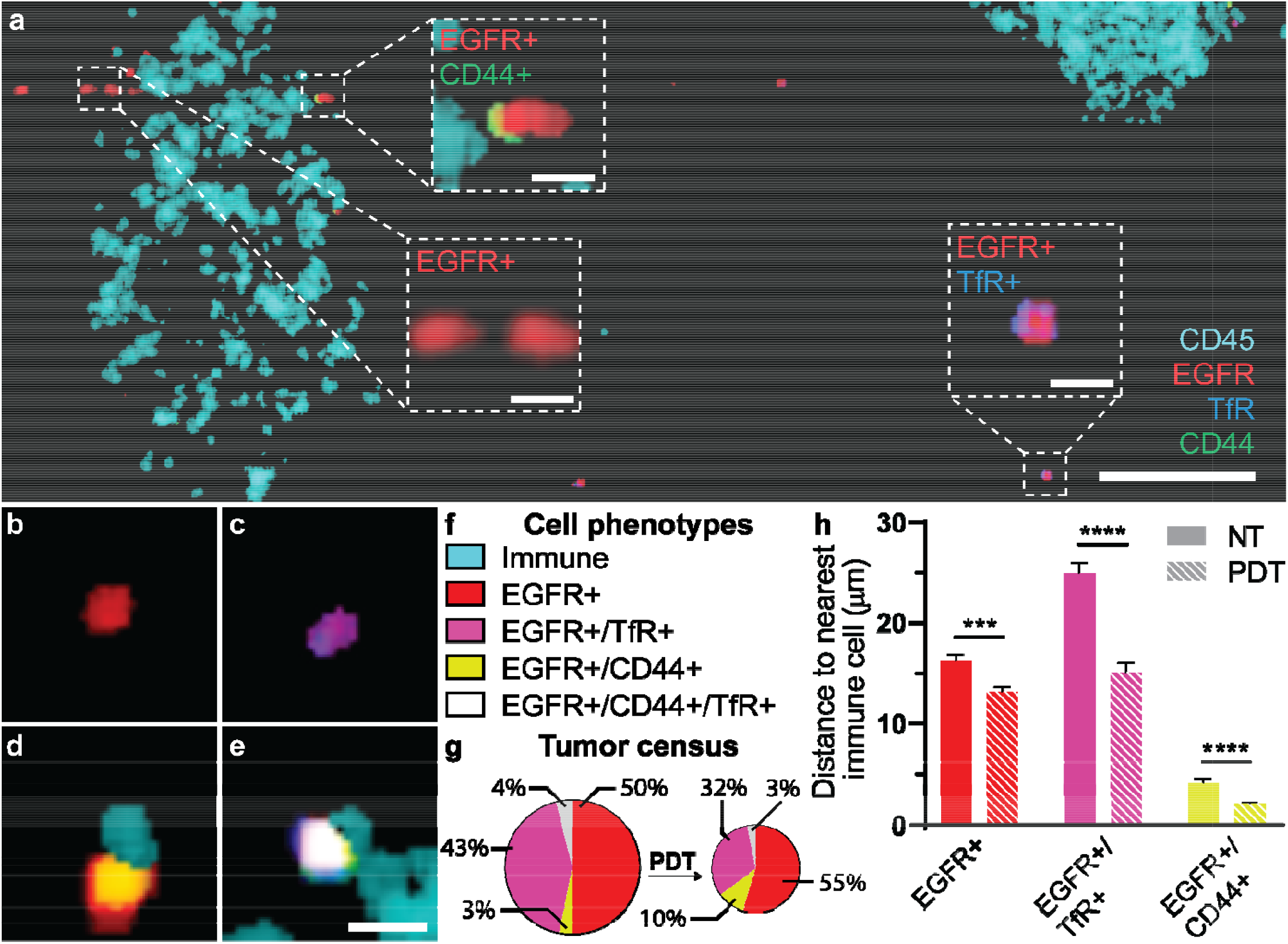
SMIRC phenotypic identification and spatial correlation. **a**, Phenotypic map of ovarian cancer from the untreated peritoneal wall of a live mouse. Scale bar 100 µm. Magnified insets highlight cancer cell subtypes (inset scale bar 10 µm). **b-e**, Representative examples of single cells of ovarian cancer with (**b**) low metabolism, (**c**) high metabolism, (**d**) CD44 overexpression and (**e**) with both upregulated metabolism and CD44 overexpression. Scale bar 10 µm. **f**, Color key for mixed biomarkers. **g**, Quantification of cancer subtype proportions with and without PDT. The overall number of cancer cells decrease, but CD44+ cells survive in the highest proportion. **h**, Quantification of mean distance between cancer cell subtypes and nearest immune cells revealing an enhanced association of CD44+ cells with immune cells, particularly after PDT, as shown in (**d**). Data are mean ± SE. EGFR-Epidermal growth factor receptor; TfR-Transferrin; PDT-Photodynamic therapy.

Intratumor heterogeneity over spatially distant sites has been implicated in impeding the prognostic value of biopsy and in treatment failure. However, evaluating microscale tumor heterogeneity over wide fields has not been possible with existing molecular imaging modalities. To rapidly acquire biomarker expression over large areas, we developed new spectral analysis techniques to enable micro-mosaiced imaging over wide fields (Figure 3a, Supplementary Video S1), thereby facilitating new opportunities for intraoperative detection and macroscale monitoring of tumor biomarker spatial fluctuations. While some features are visually apparent, extracting information from large, multidimensional data sets is a long-standing challenge (i.e., the curse of dimensionality). Fourier power spectrum analysis has been invoked to quantify fluctuations in other complex systems and provides insight into the nature of signaling between neighboring regions^22^. The spectral slope (Γ) is a statistical measure indicating how fluctuations scale with distance. A system of independent regions will have a Γ of zero (white noise); whereas enhanced cellular crosstalk can lead to Γ of unity or greater with the bulk of the fluctuation now occurring over larger areas (i.e., rare hot or dark spots). An analysis of the EGFR distribution yielded a Γ of 1.53±0.03 (Figs. 3b,d), approaching other complex systems with significant communication between neighboring regions^23^. In contrast, the Γ of the extracellular matrix-sequestered vascular endothelial growth factor (VEGF) niche distribution was significantly reduced (1.17±0.03; Figs. 3c,e). This indicates a relative increase in local heterogeneity and may represent an evolutionary strategy^24^ to broadly maximize VEGF gradient effects over several length scales, as both long and short distance gradients are known to have independent effects on angiogenesis^25^.

**Figure 3.**
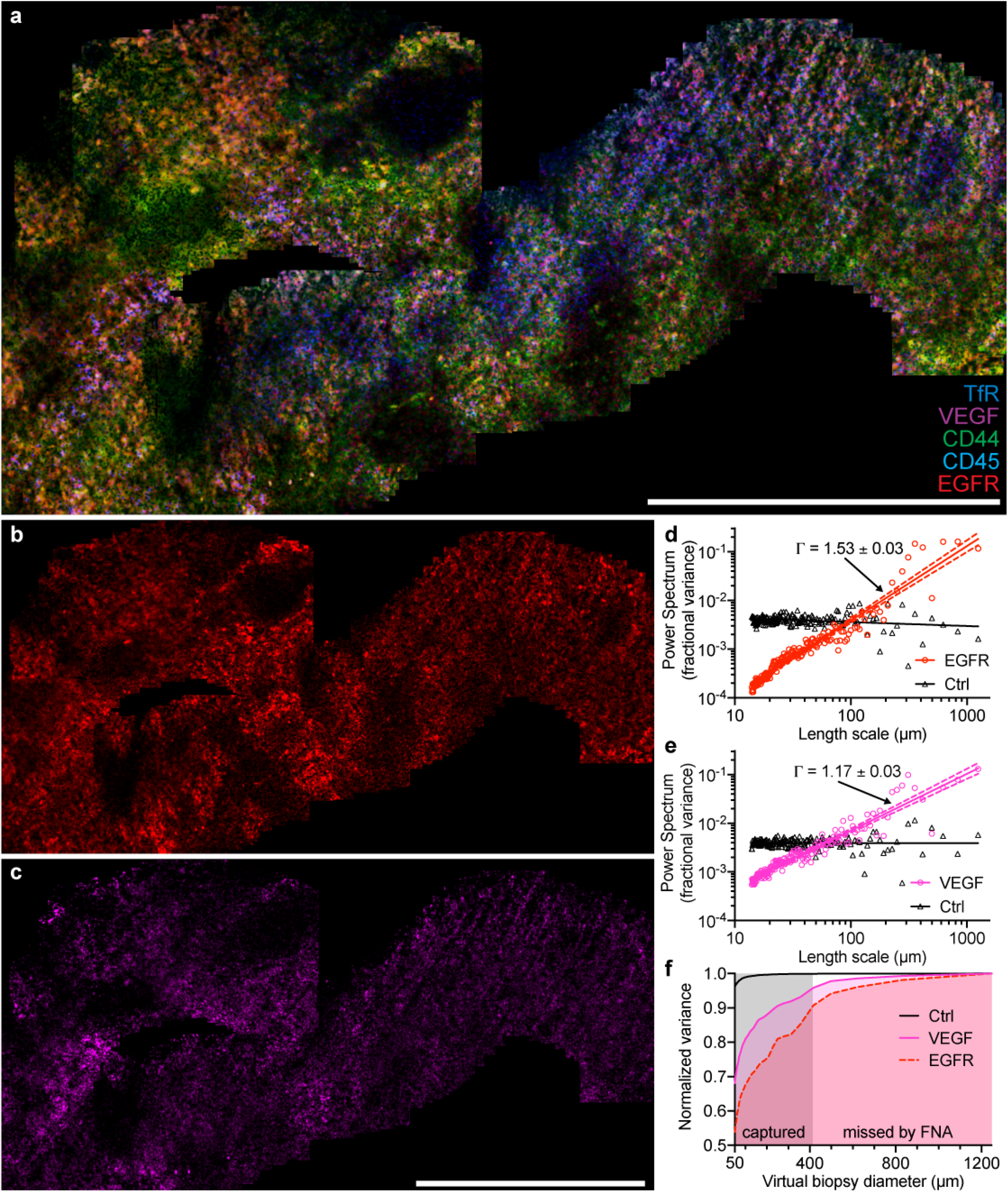
Extended field of view SMIRC with power spectral analysis of molecular heterogeneity. **a**, Mosaiced 5 target SMIRC of peritoneally disseminated ovarian cancer. **b**, The single color mosaic images of the EGFR channel and (**c**) the VEGF channel over the same region. Power spectral analysis of (**d**) the EGFR channel and (**e**) the VEGF channel quantifies the spatial dependence of the heterogeneity via the spectral slope (Γ, shown as mean ± SE). Dashed lines represent 99% CI of fit. **f**, Power spectrum calculation of heterogeneity captured over limited area quantifies the difficulty in capturing global heterogeneity with fine needle aspirate (FNA) biopsy. Scale bars 1 mm. TfR-Transferrin.

More generally, the pronounced biomarker fluctuation over large areas impedes the ability of random biopsy to capture diagnostic regions of interest^26^. While recognized as a serious limitation^1^, statistical frameworks for evaluating this phenomenon have been lacking. Here, the power spectrum provides a measure of how much heterogeneity will be captured within a randomly placed area (Figs. 3d,e). Increasing Γ indicates hotspots distributed with increasing rarity and, consequently, reduced diagnostic confidence from limited area analysis. In this tumor model, the larger EGFR Γ indicates a reduced probability to capture EGFR heterogeneity as compared to VEGF. This loss of predictive value becomes substantial as the area studied is reduced to the diameter of fine needle aspirate (FNA) sampling and below (Fig. 3f). In the future, mitigating the confounding effects of heterogeneity on *ex vivo* pathology should be possible using real-time molecular imaging technologies, such as SMIRC, to guide biopsy selection.

## Discussion

SMIRC is well placed to bridge the void between whole body radiological methods and microscale modalities such as *ex vivo* biopsy. Through the development of new optical instrumentation, mathematical models and GPU-accelerated algorithms, SMIRC overcomes the real-time requirement of *in vivo* imaging, allowing spatial correlations between different cellular phenotypes and extracellular components to be mapped and quantified with microscopic detail deep *in vivo*. As these real-time capabilities complement the macroscale visual and tactile cues currently used by surgeons for determining tumor margin^27^, SMIRC is well positioned to provide intraoperative guidance^28^. Furthermore, as the fluorescent conjugate concentrations used here are roughly 1% the therapeutic dose of other human antibody treatments, SMIRC can be tuned to have minimal biological effects^29^. The sub-millimeter diameter probe implemented here is suitable for use as is within the working channel of commercial endoscopes or, for larger field of view applications, can be scaled up with the use of larger fiber bundles or micro-mosaicking.

Collectively, these results demonstrate that longitudinal cancer cell phenotype and tumor microenvironment imaging in the context of therapeutic intervention is possible with SMIRC. This unique ability to capture cancer growth and treatment effects at the single cell level over extended areas *in vivo* and to identify problematic niches opens new possibilities for evaluating the dynamic response of existing therapeutic approaches and the rational design of combination regimens.

## Methods

### SMIRC design and implementation

The low photon counts occurring with *in vivo* multiplexed imaging precluded conventional high spectral resolution acquisition, as prescribed by the Shannon-Nyquist sampling theorem (applied in spectral space, not time), due to the prohibitively long exposure times required for accurate spectral reconstruction. However, by using prior knowledge of the spectral properties, the sampling theorem can be circumvented by finding a sparse measurement basis (i.e., configuration) from which to allow more efficient spectral estimation^10^. Basis optimization was complicated by the signal dependent Poisson shot noise, which was resolved using Monte Carlo methods (see **Supplementary Methods**). Optimal bases were systematically examined using different combinations of fluorophores, excitation conditions and frame rates using MATLAB (The MathWorks, Natick, Massachusetts, USA) implemented on a Linux cluster (Supplementary Data Fig. 1). Zemax optical (Zemax LLC; Kirkland, Washington, USA) simulations performed in parallel (Supplementary Data Fig. 5a–c) optimized for chromatic aberration and provided optical parameters for the Monte Carlo calculations. These calculations then informed the implemented optical design (Supplementary Data Fig. 2a) where the output of a 636 nm diode laser (a special order OBIS 636 nm LX 100 mW laser system with a fiber pigtail; Coherent, Wilsonville, OR) was raster scanned over the active area of the distal end of a 0.35 NA coherent fiber-optic bundle (0.95-mm-diameter FIGH-30-850N Fujikura Image Fiber; Myriad Fiber Imaging Technology, Inc.; Dudley, Massachusetts, USA) using a 10× 0.45 NA plan-apochromat microscope objective (Carl Zeiss Microscopy, LLC; Thornwood, New York, USA) with 0.7 mW of delivered power. Raster scanning was implemented with a 36-facet polygonal mirror (DT-36250-020; Lincoln Laser; Phoenix, Arizona, USA) for the fast x-axis scan, and a galvometric mirror (QS-7; Nutfield; Hudson, New Hampshire, USA) for the slow y-axis scan. Programmable hardware for horizontal (x-axis) and vertical (y-axis) scanning mirror synchronization with the data stream acquisition enabled flexible frame dimensions and image reconstruction (see **Supplementary Methods)**. The scanned laser beam is injected into the individual multimode cores composing the fiber bundle, which serve as light conduits to and from the tissue. This single laser line was used to excite the panel of targeted fluorophore conjugates. The fiber cores serve as the confocal illumination and detection pinholes to provide depth sectioning. A custom-built detector—using a 16-channel linear array photomultiplier tube with an extended sensitivity for wavelengths up to 920 nm modified with a 20 MHz amplifier (H01515M-20, Hamamatsu; Shimokanzo, Japan), and outfitted with a software-controlled, tunable Czerny-Turner spectrograph (Acton SP-2156 Imaging Spectrograph; Princeton Instruments; Acton, Massachusetts, USA)—was adjusted to collect a spectral range of 642–807 nm in approximately 11 nm steps with single photon sensitivity. A 100 µm pinhole at the spectrograph entrance reduces background signal contamination. Dichroic (LPD01-633RU-25) and notch filters (NF03-658E-25) were purchased from Semrock (Rochester, New York, USA). Four PCI express (PCIe) frame grabbers (Solios eA; Matrox Electronic Systems Ltd.; Dorval, Quebec, Canada), each equipped with 4 input channels, digitize the 16 channels of data in parallel to enable real-time frame rates.

To unmix the molecular components from the data streams at real-time speeds, a parallelized GPU implementation of non-negative least squares fitting was developed which accelerated analysis 7000 times over conventional CPU processing. To position sequentially acquired frames into a mosaic for wide-field studies, cross-correlation analysis was used to calculate the spatial shift between neighboring frames. Enhanced accuracy was obtained by using the unmixed frames, which effectively denoised the signal by making use of the biological structure to determine relevant features on which to base the frame shift. Cuts across frames were made using a minimal gradient path^30^. These methods are described in the **Supplementary Methods**.

### *In-vivo* mouse models and treatment

To mimic human peritoneal carcinomatosis, we used a mouse model previously developed in our laboratory^31^. Female Swiss nude mice, 4–6 weeks old, were injected intraperitoneally with cultured OVCAR5 cells (Fox Chase Cancer Center) at an initial inoculum of 3.15 × 10^7^ cells in single-cell suspension in PBS. To study orthotopic pancreatic cancer, experiments were carried out on 6-week-old male Swiss nude mice implanted by injection of a 50 μL volume containing 10^6^ AsPC1 cells (ATCC CRL-1682) in a 1:1 mixture of Matrigel (BD Biosciences) and culture media, as previously performed^32^. 1 hour prior to imaging, contrast agents were injected IP, as described in the **Supplementary Methods**. To perform minimally-invasive imaging, a 1 mm diameter catheter was inserted into the peritoneal cavity and the microendoscope gently pressed against the tissues of interest. The incision was sutured with a single stitch post-imaging. These mice could be imaged multiple times over several days with no apparent ill-effects. For open cavity imaging, the peritoneal cavity was surgically exposed and the microendoscope gently placed over tissue of interest. For mosaic generation, the microendoscope was glided over the tissue. To perform PDT^21^, verteporfin was administered IP at 0.25 mg/kg. After a 90 minute interval, 690 nm laser light was administered through a catheter at 90 mW/cm for 55 seconds (50 Joules). All imaged and treated mice were anesthetized with isoflurane (3.5% in 100% oxygen), and the work was approved by the Massachusetts General Hospital Institutional Animal Care and Use Committee.

### Statistical Analysis

Significance was determined with an unpaired, two-tailed Student’s *t*-test. We denoted statistical significance on the plots by asterisks with ‘***’ indicating *P* ≤ 0.001 and ‘****’ indicating *P* ≤ 0.0001. For Fig. 2h, data were taken over all cells in 9 untreated images and 10 treated images.

## Acknowledgements

We thank Clemens Alt, Brett Bouma, Pilhan Kim, Anne-Laure Bulin, Mans Broekgaarden, William Farinelli, Jerrin Kuriakose, Girgis Obaid, Seok Hyun Yun, Guillermo Tearney, and Robb Webb for assistance, as well as. Jie (Jenny) Zhao, Jermaine Henderson, Lingxian Wu, and Neema Kumar of the Photopathology Core at the Wellman Center for Photomedicine for help with *ex vivo* immunofluorescence and histopathology. This work was supported by National Institutes of Health grants R01 CA156177 (to T.H.), R01-CA160998 (to T.H.), R01-CA158415 (to T.H.), U54-EB015403 (to A.P. and T.H.), P01-CA084203 (to T.H.), F32-CA144210 (to B.Q.S.), K22-CA181611 (to B.Q.S.), and R01-CA226855 (to B.Q.S.); and the Richard and Susan Smith Family Foundation (Newton, Massachusetts) Smith Family Award for Excellence in Biomedical Research (to B.Q.S.).

## Author Contributions

B.Q.S., A.P., and T.H. conceived SMIRC. A.P., B.Q.S. and J.S. designed the optical setup. A.P. and B.Q.S implemented the opto-mechanics. A.P., M.A.S. and Z.M. designed and performed animal experiments. E.K. developed the parallelized unmixing software. R.L. developed the micro-mosaicking software. A.P. and R.L. developed the power spectral analysis. M.A.S. performed chemical synthesis of the antibody–fluorophore and – ligand conjugates. R.C.H. performed the cell counting analyses. T.H. and B.Q.S. supervised the project. B.Q.S., A.P., E.K., M.A.S., R.L., R.C.H. and T.H. wrote the manuscript. All authors contributed to editing the final manuscript.

## Notes

### Competing Interest Statement

The authors have declared no competing interest.

