## Supplemental Information for "Sparse-spectral microendoscopy for real-time visualization of tumor cell phenotype and microenvironment spatial heterogeneity *in vivo*"

**Supplementary Data Figures**
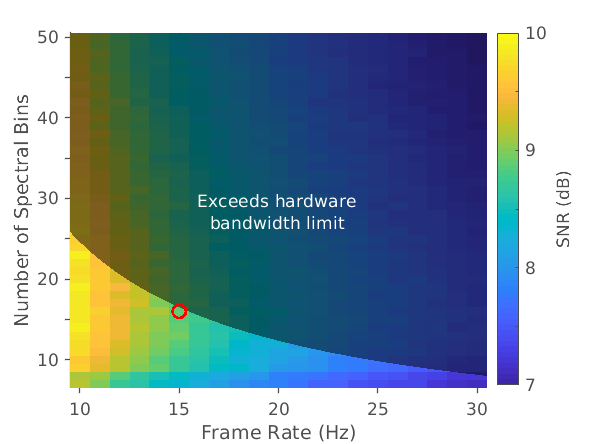

**Supplementary Data Figure 1⎪ Sparse measurement basis determination.** Spectral measurements were simulated to highlight the tradeoffs between imaging speed and spectral resolution. Monte Carlo calculations modeled spectral unmixing in the photon-starved regime influenced by illumination conditions and optical collection efficiency. These points were informed by Zemax modeling. Increasing frame rate or the spectral wavelength resolution reduced the number of photons per wavelength bin and increased the photon shot noise. Reducing spectral resolution blurred out spectral features and degraded the spectral unmixing. The combination of these effects led to the choice of measurement conditions with acceptable frame rate. The shaded area demarcates data acquisition rates larger than the maximum ability of the computer to transfer data to memory and constrained the target design regime (red circle). The simulated wavelength bins are spread out over 650 nm to 800 nm. Each point in the figure is calculated from 10,000 iterations. Alexa Fluor 633, 647, 660, 680 and 700 fluorophores were combined at concentrations designed to equalize their fluorescent contributions *in vivo* (~1 µM) for the calculation shown here.

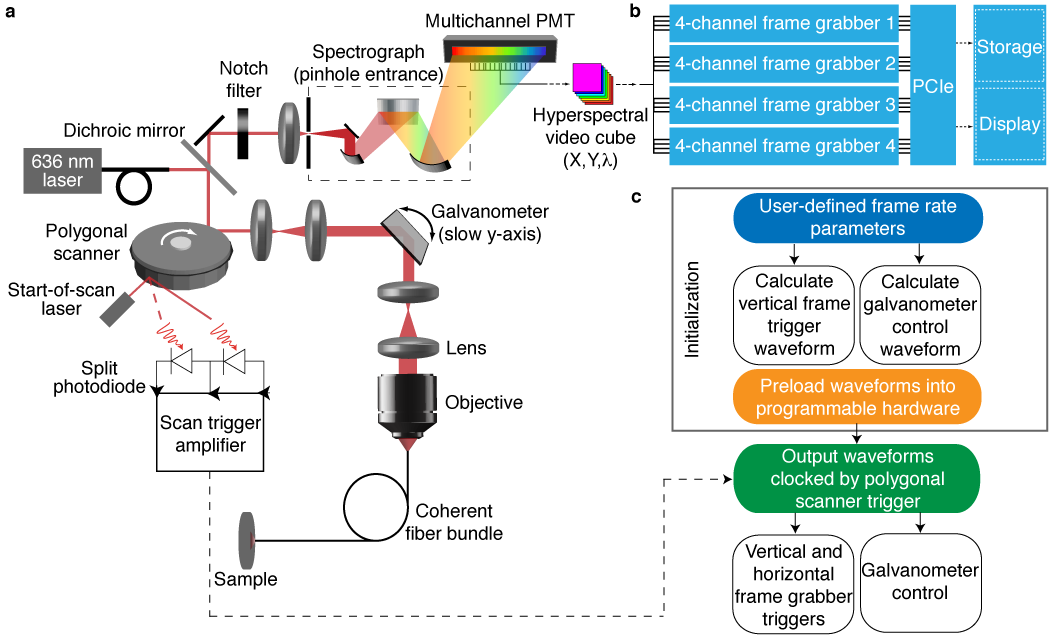

**Supplementary Data Figure 2⎪ SMIRC layout.** **a,** The opto-mechanics consists of a fiber coupled diode laser beam that is expanded and scanned over the back aperture of an objective, which couples the light into a coherent fiber bundle. The collected fluorescence is de-scanned and focused through a pinhole to remove background light. A notch filter is used to attenuate the excitation wavelength, and the remaining light passes through a spectrograph. The spectrally dispersed light is then collected by a multichannel photomultiplier tube (PMT). To detect the start of the polygonal mirror x-scan, a second off-optical axis laser is detected by a split photodiode. The resulting start of scan (SOS) signal is run through a high slew rate hysteretic comparator to remove noise before being sent to the scan engine to time the galvometric mirror and frame grabbers. **b,** The fluorescence data stream is captured by a set of multichannel frame grabbers. The data stream is directly routed to memory and display components via the Peripheral Component Interconnect Express (PCIe) bus. **c,** To synchronize the polygonal x-scan with the galvometric mirror y-scan and the image formation, a scan engine receives the SOS timing pulse and generates an analog waveform which synchronizes the motion of the galvometric y-scan. Simultaneously, a series of digital pulses is sent to the frame grabber initiating data acquisition and providing spatial information for the image cube reconstruction. Uniquely, the scan engine is initialized at run-time, allowing the scan parameters to be changed on the fly.

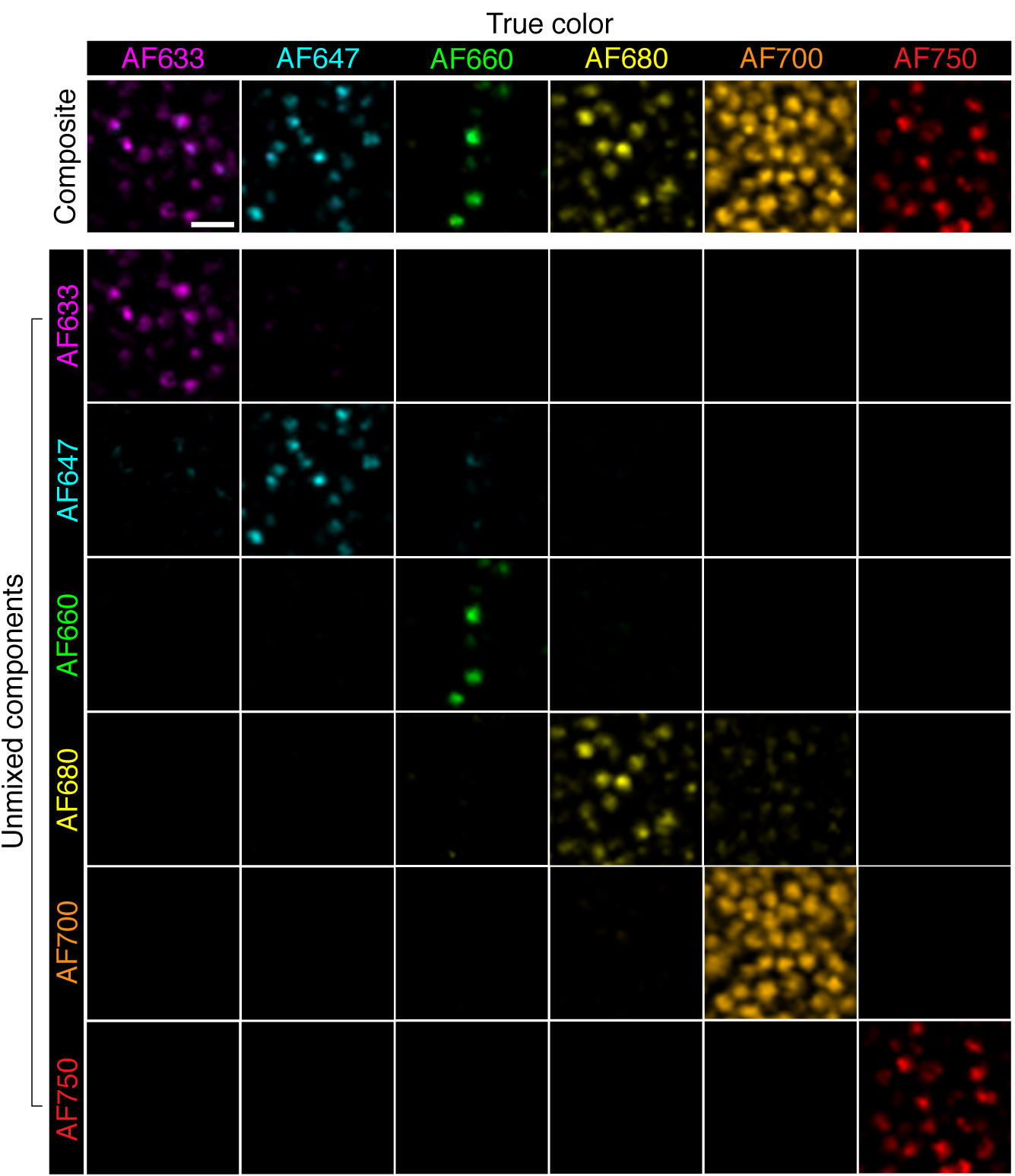

**Supplementary Data Figure 3 ⎪ Single cell *in-vitro* validation.** Fixed OVCAR5 cells were directly conjugated to Alexa Fluor (AF) dyes with NHS-ester coupling and imaged with SMIRC. 6 different conjugated cell groups were created, listed in the top title row. The composite images are the summation of all photons collected by the system. In the matrix below, the composite images are unmixed, with each row indicating the unmixed contribution of the respective fluorophore (listed on the left title bar). Diagonal elements indicate correct unmixing analysis. Each image was taken with a 60 ms exposure. 20 µm scale bar.

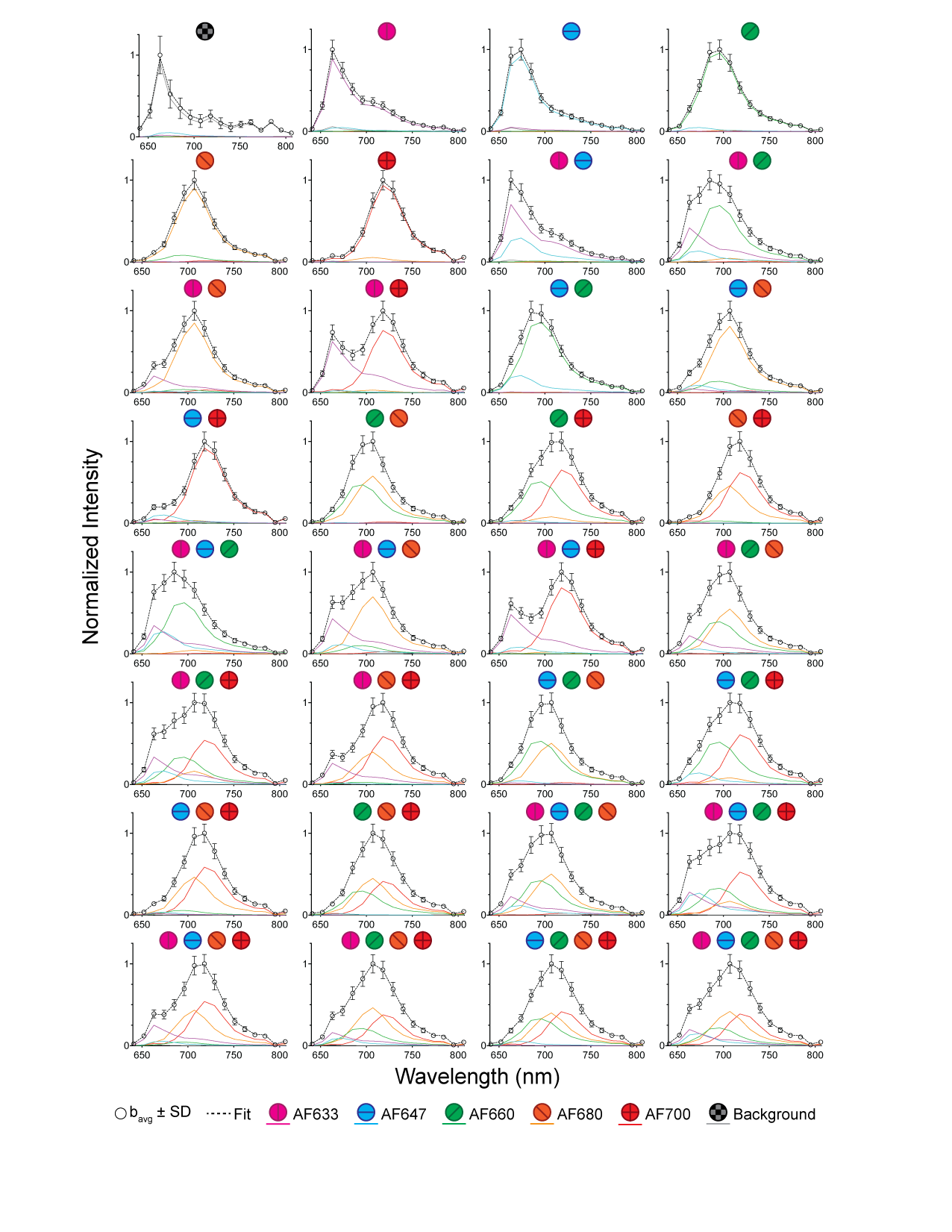

**Supplementary Data Figure 4 ⎪ Spectral unmixing of 5 fluorophores in solution.** The 32 possible permutations of 5 Alexa Fluor (AF) dyes were combined at concentrations of 0.1–10 μM and imaged with SMIRC. The raw spectra with the unmixed fluorophore spectra are shown, with each plot representing a unique fluorophore combination indicated by symbols at the top of each plot. Each fit was computed from a single 60 ms exposure. Error bars are standard deviation.

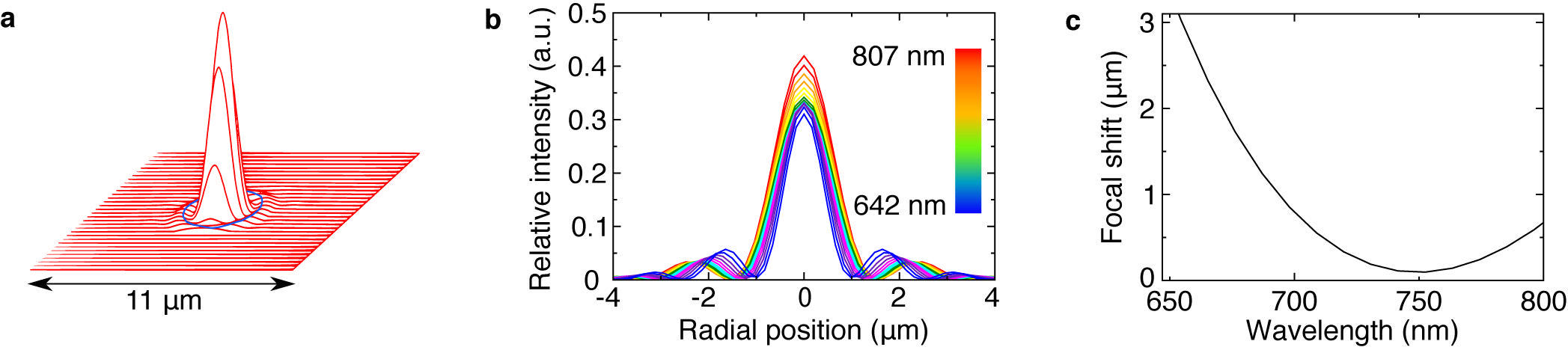

**Supplementary Data Figure 5 ⎪ Achromatic performance.** Zemax modeling of the optical design provided input for the optimized implementation. **a,** Intensity profile of the excitation beam intensity at the fiber bundle. The beam spot size is smaller than an imaging fiber core size (blue circle), indicating efficient laser coupling. **b,** Radial intensity of the emission at the pinhole entrance to the spectrometer. Chromatic aberration over a large wavelength range can lead to skewed photon collection between fluorophores with different emission profiles. Design simulations optimized the collected photons from an individual fiber to within a few microns and leads to negligible chromatic aberration effects. **c,** The focal shift as a function of wavelength at the pinhole demonstrating wide spectrum confocality.

**Supplementary Methods**

**Measurement Basis Optimization**

The pixel spectrum without measurement noise is described as a 1D vector ($\boldsymbol{S}$) and expanded in an orthonormal basis $\Psi=[\boldsymbol{\psi}_{1}\boldsymbol{\psi}_{\boldsymbol{2}}\cdots\boldsymbol{\psi}_{N}]$ as follows:

$$\boldsymbol{S}=\sum_{i=1}^{N} x_{i}\boldsymbol{\psi}_{i}$$

where the coefficient sequence $x$ is obtained from the spectrum inner product, $x_{i}=\left\langle\boldsymbol{S},\boldsymbol{\psi}_{i} \right\rangle$. The first M terms of the sequence are defined as the “sensing” basis, and the remaining terms “non-sensing”:

$$\boldsymbol{S}=\sum_{i=1}^{M} x_{i}\boldsymbol{\psi}_{i} +\sum_{j=M-N+1}^{N} x_{j}\boldsymbol{\psi}_{j}$$

An optimal sensing basis minimizes the error between the spectrum and the sensing basis reconstruction:

$$\min\left| \boldsymbol{S-}\sum_{i=1}^{M} x_{i}\boldsymbol{\psi}_{i} \right|$$

with increasing M improving the spectral reconstruction. With a suitably sparse basis, the spectral reconstruction from the sensing basis can then be unmixed using conventional non-negative least squares fitting (NNLS). This framework becomes more complicated in the *in vivo* photon starved regime due to Poissonian shot noise ($\epsilon_{P}$) which has a non-linear dependence on the photon count ($\Delta t\left\langle\boldsymbol{S},\boldsymbol{\psi}_{i} \right\rangle$). To mitigate the disruption of the shot noise on the spectral reconstruction and unmixing, an extra term is added to the minimization:

$$\mathbf{min}\left| \Delta t\sum_{i=1}^{M} x_{i}\boldsymbol{\psi}_{i}+\sum_{i=1}^{M} \epsilon_{P}\left( \Delta t\left\langle\boldsymbol{S},\boldsymbol{\psi}_{i} \right\rangle\right)-\sum_{k=1}^{5} \beta_{k}\boldsymbol{\Phi}_{k} \right|$$

where $\Delta t$ is the exposure time for pixel spectral measurement, $\boldsymbol{\Phi}$ is the fluorophore emission spectra and $\beta$ is the unmixed fluorophore abundance. The above equation is written to make explicit the dependence of the shot noise on the choice of sensing basis. Unlike the noise-free situation, increasing M does not necessarily improve the minimization, as the non-linear dependence of the shot noise on signal can dominate the other terms. The minimization is non-trivial to solve analytically, motivating the use of Monte Carlo techniques for the inclusion of the noise term followed by NNLS for the spectral unmixing. 10^4^ iterations were performed for each choice of measurement basis and frame rate to determine mean square error.

**Scan Engine**

To relink the data streams with its spatial image coordinates, a timing laser system which detects the start of the polygonal line scan (Lincoln Laser BMC-7; 10,000 rpm, 36 facet mirror) with sub-µsec precision was used to synchronize the galvometric mirror scan (Nutfield Technology QS-7). As the polygon rotated, the appearance of a new polygonal mirror facet was detected with a split photodiode (Advance Photonix Inc SD 113-2421-021) coupled to a wide-band current feedback operational amplifier in an inverting gain configuration (Analog Devices AD811) and inverting electronic Schmitt triggers (Texas Instrument CD74HC14) for backend TTL pulse generation. The timing of these scan events is critical and must be implemented in hardware to avoid image jitter. This is typically done using hardwired digital electronics for vertical and horizontal synchronization pulse trigger generation^33^. However, the resulting data acquisition rates were substantial and placed the instrumentation computer on the edge of stability. To optimize the data transfer rates and ensure stable computational operation required multiple iterations of the timing hardware, which was prohibitively time consuming. Standard externally clocked solutions were more easily modified but had unacceptable jitter due to the high acquisition clock speeds. To accelerate the optimization, a new programmable hardware implementation was developed (Extended Data Fig. 2c). The galvometric mirror and frame grabber trigger waveforms were loaded at runtime into the hardware FIFO buffer queue of a waveform generator (National Instruments PCI 6259) with simultaneous analog/digital output capabilities, and the subsequent waveform output clocked by the start of polygonal scan laser trigger. These synthesized waveforms were also used to trigger synchronized burst acquisition of the frame grabbers (Matrox Solios eA/XA). This implementation provided jitter-free synchronization of the raster scanning system with the image acquisition and enabled run-time raster scan parameter flexibility while obviating the problems associated with externally clocked trigger waveform generation. Furthermore, this design allows enhanced dynamic range by allowing a trade-off between speed and photon collection efficiency not available in conventional hardwired circuit topologies^33^.

**GPU-Accelerated Image Processing**

Each chromophore was measured independently to populate a basis spectra library containing all dyes used throughout imaging. A dark image of phosphate buffered saline was also imaged to characterize the instrument dark spectrum. Laser reflection and dark spectra were treated as additional chromophores^34^. Each pixel is assumed to be a linear combination of the basis spectra library and is therefore unmixed via linear least squares regression with a non-negative physicality constraint on the solution^35^. The non-negative least squares (NNLS) algorithm^36^ comes with significant computational cost, making real-time analysis impractical with currently available implementations^37–39^. A parallel graphics processing units (GPUs) enabled system was developed to implement faster-than-real-time NNLS unmixing, resulting in a 7,000-fold increase in unmixing speed over conventional methods. Software for GPU-NNLS was developed using the Compute Unified Device Architecture (CUDA) language and computations were carried out using one GTX 1080Ti GPU (NVIDIA). Images were normalized to the excitation laser power, and saturated pixels above the 10-bit detector limit were excluded from further analysis. A 6×6 pixel gaussian low-pass filter (σ = 4) was applied to remove fiber-bundle cores from the image. Images were then unmixed using GPU-NNLS unmixing with the user defined basis spectra library at an average rate of 9 ms/frame.

**Mosaicking**

To stitch sequentially acquired images into a widefield mosaic, the overlapping area between sequential images was analyzed using a multi-channel adaptation of micro-image mosaicking^40^. The normalized cross-correlation map is computed for each unmixed molecular abundance map in a sequential pair of images. As described in Bedard *et al.*^40^, the location of the peak of each map signifies the most probable translational correction that compensates video motion – however, there is no guarantee that the peak locations across the several imaged channels will be consistent. To overcome this and preserve spatial consistency, the cross-correlation maps of all the unmixed channels are averaged, and the location of the average peak taken as the best estimated correction for the entire image stack. Mosaics are produced by aligning frames through these corrections and implementing a graph-cut based stitching method to merge the overlapping area in a manner that minimizes edge artifacts and image disruption^30,41^.

**Phenotype analysis**

To enable facile biological interpretation of the SMIRC data sets a phenotypic mapping analysis was performed. All data taken in a mouse over a single imaging session were first thresholded with a global isodata algorithm^42^. All sub-threshold pixels were then set to zero. This was performed on each unmixed channel. The remaining unmixed pixels were checked for overlap with threshold positive CD45. Overlapped pixels were also set to zero. The Euclidian distances between these remaining cancer cell subtype pixels to the nearest immune pixels were analyzed for subtype niches and treatment dependence. Due to the large dynamic range of the data, faint pixels could not be visualized against bright pixels on a linear color brightness scale. To facilitate visualization, color brightness was remapped to a logarithmic scale, and inset figures were de-pixelated by breaking each pixel into a 3x3 grid and performing bicubic interpolation.

**Power spectral analysis**

Fourier power spectral analysis allows a framework from which to investigate the spatial dependence of heterogeneity. To perform, a 2.496 mm x 0.484 mm mosaiced rectangular area was 2D Fourier transformed using a Hamming window to mitigate spatial frequency crosstalk^43^. A fast Fourier implementation allowed real-time analysis of the unmixed channels. The square magnitude of the transform was then radially averaged and plotted as a function of spatial period to extract the spectral slope. Using Parseval’s theorem, which equates the integral of the power spectrum over the entire transformed space to the variance (a measure of heterogeneity), the spatial frequencies of interest were integrated over to provide an estimate of captured variance as a function of analyzed area. Controls were virtually created by randomly scrambling the unmixed images and repeating the calculations.

**Fluorescent contrast agents**

Antibodies were conjugated with NHS-ester chemistry to Alexa Fluor dyes (Thermo-Fisher) for 2 hours at room temperature before size-exclusion column purification. This reaction scheme led to a fluorophore/antibody ratio of 4 for anti-CD45 (Biolegend 30-F11), 8 for cetuximab (ImClone LLC), 8 for anti-CD44 (Bioxcell IM7), 4 for anti-MUC16 (Abcam X75) and 7 for bevacizumab (Roche). Peanut agglutinin (Thermo-Fisher L32460) and transferrin (Thermo-Fisher T23362) were purchased pre-conjugated to Alexa-Fluor dyes. The injected masses of conjugate used here were 31 µg anti-CD45, 12.5 µg cetuximab, 5 µg anti-CD44, 25 µg anti-MUC16, 13 µg bevacizumab, 1 µg peanut agglutinin and 20 µg transferrin. These were injected IP in 1 ml PBS.

**Specific experiment details**

Orthotopic pancreatic minimally invasive imaging was performed 43 days post-inoculation. After insertion of the catheter, the microendoscope was manually guided toward the tumor, which was externally palpable by touch. Strong fluorescence was clearly apparent when imaging the pancreatic tumor, as opposed to other tissues which were dim. Peritoneal carcinomatosis imaging was performed 9 days (Fig. 2) and 20 days (Fig. 3) post-inoculation. These time points were chosen to demonstrate imaging of early and late-stage dissemination. When examining treatment effects, PDT was performed on day 8. To provide an internal control, PDT was performed on the region of the peritoneal cavity near the right hind leg only. The left hind leg region was used as an untreated comparison. For mosaic imaging of peritoneal carcinomatosis, the peritoneal cavity was surgically exposed and visible, mimicking open peritoneal surgery. The microendoscope was glided over a section of the peritoneal wall for several seconds and images captured at 16.7 fps.

**Supplementary Video S1. ⎪ Online SMIRC video and mosaicking.** 5 channel live animal SMIRC imaging of a peritoneal carcinomatosis mouse model was performed at 17 fps and is shown here at the same frame rate. The right panel displays the SMIRC field of view (scale bar 100 µm). The left panel displays the widefield mosaic formation as images are acquired (scale bar 500 µm). Targets imaged: transferrin (dark blue), VEGF (magenta), CD44 (light green), CD45 (cyan) and EGFR (red).
